# Stimulus dependent modulation of perceptual filling-in is predicted by the properties of early visual cortex

**DOI:** 10.64898/2026.07.01.730966

**Authors:** Anna Razafindrahaba, Kenshu Koiso, Vincent van de Ven, Federico de Martino, Peter De Weerd, Mark J. Roberts

## Abstract

Filling-in occurs during the perceptual disappearance of a blank figure presented on a textured background. Current models of perceptual filling-in are based on a two-stage model where the figure boundary weakens after a period of adaptation, followed by the spreading of the background representation into the region representing the figure. This suggests a competition between figure boundary and background representations whereby filling-in is facilitated by a weaker boundary representation and a stronger background representation. Here, we test this interpretation, by using the oblique effect and surround-modulation suppression, which are functional properties of early visual cortex that modulate the expected strengths of the responses to the background texture and to the figure boundary. In a sample of *N=58* participants, we found more filling-in with background textures of cardinal compared to oblique orientations (earlier onset time, with more and longer episodes of filling-in per trial), in line with a known, stronger neuronal response for cardinal than for oblique orientation in early visual cortex. We found more filling-in when the main axis of the rectangular figure was iso-oriented rather than cross-oriented with the background texture (more and longer episodes of filling-in per trial, but no change in onset time), in line with a lower response to oriented stimuli when surrounded by iso-oriented flankers compared to cross-oriented flankers. Overall, our results support the two-stage model and suggest the involvement of early visual cortical areas characterized by the oblique effect and orientation-tuned surround-suppression.

## Introduction

Filling-in is a perceptual phenomenon in which information is perceptually interpolated across areas where input is missing. In Troxler’s stimulus (Troxler, 1804), a grey figure presented away from the center of gaze becomes filled-in by the surrounding line texture background after prolonged fixation, including a period of reduced microsaccades (Troncoso et al., 2008). In the Troxler paradigm, the extent to which the figure boundary is stabilized on the retina and the ensuing adaptation and weakening of the boundary representation are thought to be preconditions for perceptual filling-in to occur (Gerrits et al., 1966; Gerrits & Vendrik, 1970; Walls, 1954). Following sufficient adaptation of the neural representation of the figure boundary, information from regions representing the background may be interpolated into the region representing the figure, supported by horizontal connections in low-level visual cortex and/or feedback from high-level visual cortex (De Weerd et al., 1995; Dennett, 1992; Spillmann & De Weerd, 2003). Thus, in the Troxler stimulus, perception of the figure versus perception of filling-in may represent a competition between the representation of the background and of the figure boundary, with filling-in resulting from either a stronger representation of the background, or a weaker representation of the boundary. Here, we tested this interpretation using a stimulus that consisted of a grey rectangle placed on a background texture of oriented line elements to take advantage of two known tuning properties of early visual cortex. The “oblique effect” (Furmanski & Engel, 2000; Mikellidou et al., 2015) allowed us to manipulate the expected strength of the background representation, and orientation-tuned surround suppression (Chen & Tyler, 2001; Maniglia et al., 2022) allowed us to manipulate the expected strength of boundary representation. We hypothesized that filling-in would be most prevalent for stimulus configurations characterized by a strong background response and a weak boundary response.

With respect to the background, we hypothesized that cardinal orientations in the background texture would elicit more filling-in than oblique textures, in line with the stronger population response to cardinal orientations in early visual cortex (Furmanski & Engel, 2000; Hubel & Wiesel, 1962, 1968; Self et al., 2016; Sun et al., 2013). The early visual cortex in animal models is characterized by orientation-tuned neurons (Hubel & Wiesel, 1962, 1968). This finding has been replicated in the human early visual cortex with studies using multi-unit electrophysiology recordings, or magnetic resonance imaging (Patten et al., 2017; Self et al., 2016; Shen et al., 2014; Sun et al., 2013; Yacoub et al., 2008).Yet, not all orientations are represented equally. In animal models such as the cat and the monkey, a higher proportion of cells are tuned to cardinal orientations than to obliques (Hubel & Wiesel, 1968; Li et al., 2003; Pettigrew et al., 1968; Shen et al., 2014). Likewise, in humans, functional magnetic resonance studies have found more voxels tuned to cardinal than oblique orientations, suggesting stronger or more reliable neural activity for cardinal than oblique orientations (Furmanski & Engel, 2000; Sun et al., 2013). The lower population response for oblique compared to cardinal orientations has been termed the “oblique effect” (Appelle, 1972). This effect offers an interesting avenue for modulating the level of population activity in primary visual cortex. Because we can anticipate a higher population response for texture backgrounds of cardinally rather than obliquely oriented elements, we expected backgrounds composed of cardinally oriented line elements to facilitate perceptual filling-in.

In addition to the oblique effect, some studies have reported differences between vertical and horizontal orientations, and between obliques in both neural responses and visual performance (Alink et al., 2017a; Hansen & Essock, 2004; Leventhal & Schall, 1983; Li et al., 2003; Ling et al., 2015; Maloney & Clifford, 2015; Mannion et al., 2010b, 2010a; Menceloglu et al., 2023; Sasaki et al., 2006; Schall et al., 1986; Westheimer, 2003, 2005; Yacoub et al., 2008). fMRI studies have shown a slight overrepresentation of vertical orientations in early visual cortex, with more voxels and greater columnar regularity or density for vertical orientations (Alink et al., 2017a; Freeman et al., 2011, 2013; Hansen & Essock, 2004; Maloney & Clifford, 2015; Mannion et al., 2010b, 2010a; Yacoub et al., 2008). Some of these fMRI studies also reported a preference for radial orientations, pointing towards the center of gaze, over tangential orientations, pointing away from the center of gaze (Alink et al., 2017a; Freeman et al., 2013; Ling et al., 2015; Maloney & Clifford, 2015; Mannion et al., 2010a, 2010b; Sasaki et al., 2006), consistent with psychophysical (Bennett & Banks, 1991; Hong, 2015; Menceloglu et al., 2023; Rovamo et al., 1982; Temme et al., 1985; Westheimer, 2003, 2005) and physiological data (Leventhal & Schall, 1983; Levick & Thibos, 1982; Schall et al., 1986). However, opposing findings have also been reported. For example Li et al. (2003) observed more cells with a preference for horizontal than vertical orientations, while Mannion et al. (2010b) reported that the preference for radial over tangential orientations depended on the stimulus configuration. Together, these findings suggest a subtle preference for vertical over horizontal and radial over tangential stimuli, which may facilitate filling-in for those orientations. However, these differences appear smaller, more stimulus-dependent and less universally observed than for the oblique effect.

With respect to the strength of the figure boundary representation, we expected variation depending on the orientation of the rectangular figure’s length axis relative to the orientation of line elements in its background. A prominent property of orientation-selective units in early visual cortex is the modulation of their response by local orientation context, whereby the presence of flankers in the receptive field surround generally leads to suppression of the firing rate (Gilbert et al., 2000; Hubel & Wiesel, 1965; Kapadia et al., 1995; Maniglia et al., 2011; Nurminen & Angelucci, 2014; Polat & Sagi, 1993; Self et al., 2014; Yu et al., 2002). Surround-modulation is generally suppressive (Adini et al., 1997; Chen & Tyler, 2001; Gilbert et al., 2000; Maniglia et al., 2011; Nurminen & Angelucci, 2014; Polat & Norcia, 1996; Polat & Sagi, 1993; Solomon & Morgan, 2000; Yu et al., 2002) yet, the degree of suppression is influenced by the orientation difference between the central element and the surrounding flankers, with iso-oriented stimuli being the most suppressive (Henry et al., 2013; Sakai & Nishimura, 2006; Self et al., 2014; Shushruth et al., 2013; Sillito et al., 1995). In the context of Troxler filling-in, neurons representing the edge of the figure may be suppressed by the surrounding texture elements, with higher suppression expected along sections of the figure’s boundary where its orientation matches that of the local line elements in the background texture. Here, we used rectangular figures, where the longer edge, three times the length of the short edge, could be iso- or cross-oriented to the background texture, thereby modulating the expected overall level of suppression along the boundary representation in early visual cortex. Accordingly, we expected a weaker boundary representation and enhanced filling-in, for iso-oriented figures than for cross-oriented figures.

As an additional, exploratory, manipulation, we tested whether presenting the figure in the upper or lower hemifield modulates perceptual filling-in. To our knowledge, no study has directly compared filling-in between hemifields: previous work either presented figures exclusively in one hemifield, either upper (e.g., Ramachandran & Gregory, 1991; Sakaguchi, 2001; Shimojo et al., 2001) or lower (e.g., De Weerd et al., 1995, 1998), or, when both were used (e.g., Meng et al., 2005; Spillmann & Kurtenbach, 1992; Troncoso et al., 2008), did not analyze hemifield differences. This is despite evidence for a lower field advantage in the perception of illusory contours (Rubin et al., 1996; Shapley et al., 2003). Whether this asymmetry extends to perceptual filling-in remains an open question.

In summary, we tested whether manipulating the expected strengths of the background and figure boundary representations, according to the properties of early visual cortex, would modulate perceptual filling-in. We also explored whether presenting the figure in the upper or lower hemifield would modulate perceptual filling-in in line with what has been reported for other aspects of vision.

## Material and methods

### Participants

Sixty-nine healthy adults (60 females, 9 males, mean age 21.2 years, *SD=*1.68) with normal or corrected-to-normal vision participated in this study. Participants were recruited from the second-year Psychology Bachelor program at Maastricht University. Participants provided informed consent and received study credits in exchange for participation. Ethics approval was obtained from the Ethics Review Committee Psychology and Neuroscience, in accordance with the declaration of Helsinki. Participants with a self-reported history of neurological and psychiatric disorders were excluded. Seven participants were excluded due to incomplete data, 2 additional participants were excluded as their data suggested nearly continuous filling-in, which we interpreted as misunderstanding the task instruction, and 1 additional participant was excluded from further analysis due to no filling-in reports giving a final sample of 58 participants for the main dataset. For the analysis of eye-tracking data (microsaccade rate, direction and fixation), a further 4 participants were excluded due to incomplete eye-tracking data, leaving 55 participants for the eye-tracking analysis.

This sample approximates the target number of participants, calculated with a power analysis using G-Power3 (Faul et al., 2007), based on a 4×2 within-subjects ANOVA corresponding to the main hypotheses of the study. Main effects of power 0.8 and medium size effect of 0.25 required 30 participants for the factor ‘background representation’ and 34 for the factor ‘boundary representation’. Since we planned two groups of participants corresponding to the upper and lower visual fields, we had set the target number of participants to 68.

### Apparatus and stimuli

Participants sat with their head in a chin-and-headrest at a fixed distance of 55 cm from a computer monitor (field of view of size 38 x 30 cm, resolution 1280 × 1024 pixels, refresh rate 60Hz). The display had a size of 38.12° x 30.15° and was filled with a dynamic texture field except for a grey rectangular figure placed at the center of the screen. The dynamic texture field refreshed at 20 Hz by pseudo-randomly selecting one from 25 random patterns, while preventing immediate repetitions of the same pattern. The patterns were created by randomly setting non-overlapping white bars of size 0.72° x 0.06° (24 x 2 pixels) on a black background. The orientation of the white bars could be oblique, negative-oblique, horizontal or vertical yielding 4 background conditions for the within-participant background factor (Figure 1B). The grey rectangular figure subtended 8° x 2.5° and was equiluminant to the mean luminance of the texture (35 cd/m^2). The orientation of the rectangular figure could be vertical or horizontal yielding 2 figure orientation conditions for the within-participant figure orientation factor. Both the background and figure orientation factors yield 8 possible stimuli configurations within-participant. The fixation spot was a white spot surrounded by a green ring presented at an eccentricity of 14.58° diagonally away from the center of the screen. Its location could be at the top-right or the bottom-right corner of the screen, placing the figure in the lower or upper visual field respectively yielding two stimulus presentation conditions counterbalanced between participants.

**Figure 1:**
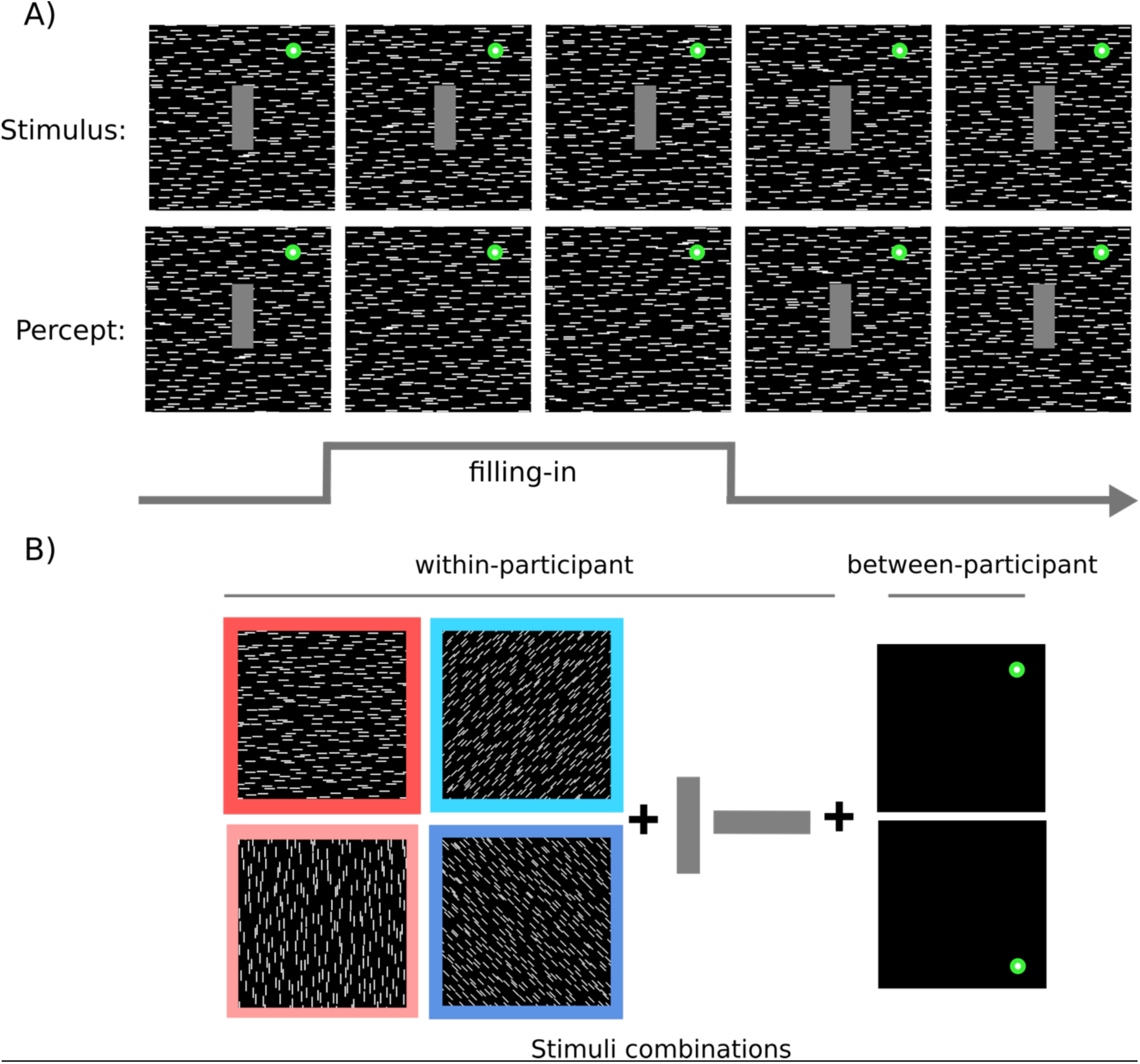
**(A)** Filling-in illustrated through the difference between the visual stimulus and the perceived stimulus. Top row: visual stimulus - Middle row: percept experienced by the participant where the central grey figure is filled-in with the surrounding texture - Bottom row: participant behavior when experiencing filling-in. **(B)** Available stimulus combinations with choices of background orientation (4), figure orientation (vertical or horizontal) (2) (as within-participants factors) and fixation spot location (2) (as between-participant factor) yield a factorial design of 4 x 2 x 2. Colored frames indicate color coding used in later figures.

### Experimental procedure

#### Perceptual Filling-in task

Participants were instructed both with a written document and verbally while viewing the stimuli on the screen prior to the task. They were instructed to maintain fixation on the fixation spot and to report full perceptual filling-in by pressing the space bar key on the keyboard and to keep pressing for as long as the percept of complete filling-in lasted.

Participants were asked to report only complete filled-in percepts and were instructed to not respond to partially filled-in percepts, and to release the space bar as soon as filling-in no longer appeared complete. The experimenter then verbally described possible configurations of partially filled-in and faded-out percepts. Specifically, participants were instructed to not respond to percepts in which the figure would darken or fade without filling-in of the texture elements across the entire figure. In addition, to set a criterion for fully filled-in percepts, participants were shown on the screen a full texture to illustrate full filling-in of the figure. We applied these detailed instructions to enforce an unambiguous response to fully filled-in percepts versus partial filling-in or perceptual fading.

In the main task, participants completed 8 blocks of 16 trials, corresponding to 2 presentations (pseudo-randomly interleaved to ensure no successive repetition of conditions per block) of each condition per block, yielding 16 total repetitions of each condition and 128 total trials. Each trial lasted 25s with an inter-stimulus interval of 1s.

#### Eye-data recording

Eye movements and pupil size of the right eye were tracked using the pupil-glint vector at a sampling rate of 225.4 Hz using the ViewPoint eye-tracker (Arrington Research, Scottsdale, AZ) and ViewPoint software version 2.9.5.128. Calibration was performed using the in-built 16-points calibration procedure at the beginning of each run, complemented by a participant-controlled recalibration task using 5 points (Nystrom et al., 2013) in which participants were instructed to fixate a fixation spot and maintain fixation while holding the space bar key until the fixation spot disappeared. Each fixation spot was shown for 1 s. This was repeated for each of the 5 locations. The recalibration task occurred at the beginning and halfway through each block of 16 trials of filling-in.

### Data analysis

#### Behavioral data analysis

##### Quantification of perceptual filling-in behavior

We first calculated the time course of filling-in behavior as a preliminary measure to quantify perceptual filling-in across time, calculated as the proportion of the participant’s reports of filling-in per time point across trials. Further, we calculated four primary measures to quantify perceptual filling-in: filling-in onset, total filling-in duration per trial, mean filling-in episode duration, and filling-in episode count. Filling-in onset was calculated as the onset time of the first filling-in episode in a trial. Total filling-in duration per trial was calculated by summing the durations of all reported instances of filling-in per trial. This measure indicates the overall amount of filling-in per trial and is a combination of the duration of each filling-in episode and the number of filling-in episodes, but it does not distinguish between, for example, several shorter episodes of filling-in vs fewer longer episodes within a trial. We therefore also calculated mean filling-in episode duration as the mean duration of each episode of filling-in as a quantification of the stability of filling-in, and the mean number of filling-in episodes per trial as the filling-in episode count.

##### Definition of conditions for factors ‘texture’ and ‘boundary’ representations

To investigate the question of texture representation, we considered only the ‘background’ factor and discarded the ‘figure orientation’ factor as a within-participant analysis. We first grouped together horizontal and vertical background orientations into the ‘cardinal’ condition and the positive and negative orientations into the ‘oblique’ condition, yielding 32 trials per condition. We next investigated differences between the four background orientations; horizontal, vertical, radial and tangential. The positive and negative-oblique orientations appear as ‘radial’ or ‘tangential’ depending on the position of the fixation position. Thus, we combined the positive-oblique condition from the lower hemifield group with the negative-oblique from the upper hemifield group as the ‘radial’ condition, and combined the remaining condition as the ‘tangential’ condition. There were 16 trials per condition. To investigate the question of boundary representation, we considered the interaction between the figure orientation and the background orientation as a within-participant analysis. For this analysis, since the orientation of the main axis of the central figure could only be vertical or horizontal, we considered only the horizontal and vertical background textures to contrast iso-oriented or cross-oriented stimuli. We grouped together the horizontal background and figure and vertical background and figure for the iso-oriented condition and horizontal background and vertical figure and vertical background and horizontal figure for the ‘cross-oriented’ condition, yielding 32 trials per condition.

### Statistical tools

Descriptive statistics, mean, standard deviation, median, and interquartile range, were calculated with the *numpy* library (Python 3.8). Inferential statistics, including both parametric and non-parametric tests, were employed for hypothesis testing. Prior to hypothesis testing, assumptions of normality and homogeneity of variance were evaluated. Normality of the data distributions was assessed using the Shapiro-Wilk test. Homogeneity of variance was examined using Levene’s test. These tests were conducted to ensure the validity of subsequent inferential statistical analyzes. For within-participant comparisons, since the data violated these assumptions, the non-parametric Wilcoxon signed-rank test or the sign test was preferred when comparing subsets with skewed distributions. Welch’s t-test was used when comparing subsets with unequal variances but still following normal distributions. Between-participant comparisons were conducted using the Mann-Whitney U-test due to the non-normal distribution of the data. For between-group comparisons ‘upper’ and ‘lower’ hemifield presentation, any non-significant differences for the Mann-Whitney-Wilcoxon U test were evaluated with Bayes Factors using a default Cauchy prior with scale 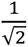. For the Bayes Factor analysis, we employed the functions in R provided by (Van Doorn et al., 2020). We used Jeffrey’s scale to interpret the evidence in favor of the null-hypothesis provided by Bayes Factors (Jeffreys, 1998). The scale suggested by Jeffreys considers Bayes Factor (*BF*_10_) values below 1/3 as moderate to strong evidence and between 1/3 and 1 as anecdotal evidence in favor of the null-hypothesis. The Friedman test was used as a non-parametric alternative to repeated-measures ANOVA for comparisons between more than two levels within a factor. We preferred the non-parametric alternative to repeated-measures ANOVA since the data violated the normality assumptions. Post-hoc tests, specifically the Wilcoxon signed-rank test with Holm-Bonferroni correction, were employed to further analyze significant findings. For temporal analyzes, multiple comparisons were corrected with a cluster-based permutation test with 1000 permutations of condition labels.

All previously mentioned statistical analyzes were performed using Python 3.8 with the *scipy.stats* library with an uncorrected (except in the temporal analysis) two-tailed significance threshold *alpha*=0.05.

#### Eye-data processing

Eye-data was processed using MATLAB R2023a. Epochs were extracted from the eye-data text files using the triggers defining start and end of the trials block per block. After matching epochs with trial information including background and figure orientation and trial number, we downsampled the data to 100Hz to reduce necessary computing power.

For microsaccades (MS) analysis, we used the Engbert and Kliegl (2003) microsaccade detection algorithm on smoothed horizontal and vertical eye signals over 7 data samples and a velocity threshold criterion of 3 standard deviations above the median velocity. We first analyzed the entirety of the trial. In the second analysis, we analyzed 10s epochs centered on participant reports of the onset of filling-in. To test for changes in microsaccade rate associated with filling-in onsets we compared the observed time course against a permuted null distribution created by randomly shuffling filling-in onset times between trials. The statistical difference between the distributions was assessed with cluster-based permutation tests. We used the same approach for filling-in episode offsets.

For the microsaccade direction analysis, we calculated the angle of each detected microsaccade, and determined distribution of microsaccade directions using 10° bins to cover the −180° to 180° range. We then converted this circular distribution to complex vectors, where each vector’s angle corresponded to a bin center and its magnitude to the proportion of microsaccades in that directional bin. To test for systematic differences in the circular distributions of vector angle between paired conditions, we computed per participant the circular mean direction per condition, and the paired circular difference. To test whether these differences deviated from a continuous distribution with a circular median of zero, we used the sign test. Similarly, we tested the difference in mean vector length between paired conditions in the sample of participants using the Wilcoxon signed-rank test.

To test for overall gaze fixation stability and for systematic differences in gaze positioning between conditions, we extracted horizontal and vertical gaze coordinates from each trial in each 16-trial testing block. These were corrected for small drifts by subtracting from them the gaze coordinates measured during the nearest preceding recalibration block, which occurred at the beginning of each testing block (preceding trial 1), and halfway through the block (preceding trial 9). We used the corrected gaze positions to calculate the distribution of gaze positions relative to the fixation spot. In addition, to assess potential gaze repositioning towards or away from the figure across stimulus conditions, we computed the distance between the gaze position and the figure. Differences in fixation biases across stimulus conditions relative to the figure location across paired conditions were assessed using the Wilcoxon signed-rank test.

## Results

### Sample analysis

The study included 58 participants with each participant completing 128 trials yielding 16 trials per stimulus configuration (4 background orientations x 2 figure orientations yielding 8 possible stimulus configurations, see Figure 1B). Across all participants and trials, the mean filling-in (FI) onset per participant (Figure 2B) was 12.20s (*SD=*2.32) into the trial. The mean total filling-in duration per trial per participant (Figure 2A) was 4.98s (*SD=*4.26). The mean duration of a filling-in episode per participant (Figure 2C) was 3.17s (*SD=*2.31). The mean number of filling-in episodes (Figure 2D) was 1.42 episodes per trial (*SD=*0.76). Figure 2 shows a skewed distribution for total filling-in duration per trial and a gaussian distribution for filling-in onset. There was no evidence suggesting a multimodal distribution for either measure. These results indicate that perceptual filling-in is a reliable phenomenon across a large sample of participants without prior experience in participating in this type of experiment.

**Figure 2:**
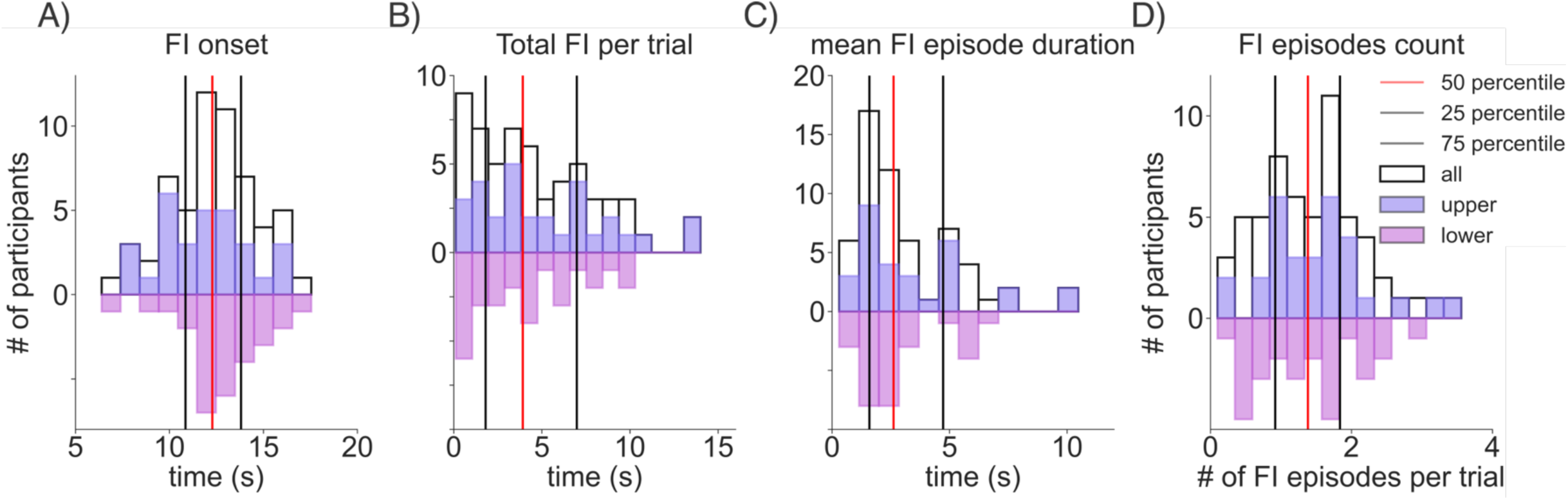
White histogram: Filling-in behavior in a large pool of participants (N=58). Colored histograms: Distribution split between the upper and lower hemifield presentation groups. Vertical lines show the first quantile (grey line), median (red line) and third quartile (grey line) of the full sample. Histograms show respectively **(A)** filling-in onset time **(B)** mean total filling-in duration per trial **(C)** mean filling-in episode duration **(D)** mean filling-in episode count per trial.

To investigate whether filling-in differed between the upper and lower visual fields, participants were divided into two groups where the figure was presented in the lower (*N*=29) or the upper (*N*=30) visual field. In the ‘lower’ group, the mean total filling-in duration per trial was 4.78s (*SD=*4.82, Figure 2B) and the mean filling-in onset was at 12.63s (*SD=*2.29, Figure 2A). In the ‘upper’ group, the mean total filling-in duration per trial was 5.18s (*SD=*3.71, Figure 2A) and the mean filling-in onset was at 11.78s (*SD=*2.31, Figure 2B). A Mann-Whitney U-test revealed no statistical differences, neither in total filling-in duration per trial (*U=*383.0, *p=*0.43, *BF*_10_=0.51) nor in filling-in onset time between the groups ‘lower’ and ‘upper’ (*U=*537.0, *p=*0.12, *BF*_10_=0.78). In the group ‘lower’, the mean filling-in episode duration was 2.76s (*SD=*1.78, Figure 2C) and the mean filling-in episode count was at 1.32 (*SD=*0.75, Figure 2D). In the group ‘upper’, the mean filling-in episode duration was 3.57s (*SD=*2.63, Figure 2C) and the mean filling-in episode count was at 1.51 (*SD=*0.75, Figure 2D). A Mann-Whitney U-test revealed no statistical differences, neither in mean filling-in episode duration (*U=*385.0, *p=*0.33, *BF*_10_=0.50) nor in filling-in episode count between the groups (*U=*380.0, *p=*0.41,*BF*_10_=0.33). Since we found no evidence for different filling-in behavior between upper and lower visual fields, while acknowledging that the Bayes Factor suggested anecdotal evidence for the null hypothesis, we combined the two groups for the following analyzes to enhance statistical power.

### A stronger texture representation elicits more filling-in

Previous studies have shown a greater population response in early visual cortex to cardinal orientations than to non-cardinal orientations (Campbell & Maffei, 1970; Furmanski & Engel, 2000; Mikellidou et al., 2015; Proverbio et al., 2002; Sun et al., 2013). We manipulated the orientation of the background’s texture elements between cardinal and oblique to assess whether modulating the strength of the expected neural response to the texture would modulate filling-in. Following our hypothesis, we first combined the horizontal and the vertical orientations into the group ‘cardinal’ and the negative and the positive-oblique orientations into the group ‘oblique’ within participants, yielding 32 trials per condition. Figure 3A shows that the proportion of filling-in reports rate increased over time for both cardinal and oblique orientations with the cardinal orientations associated with more filling-in than the oblique orientations. To ascertain the significance of this difference, we performed a Wilcoxon signed-rank test at each timepoint, with multiple comparisons corrected for by a cluster-based permutation test. This analysis revealed several temporal clusters (black horizontal bars) for the majority of the trial duration, indicating a statistically significant difference between the oblique and cardinal conditions. We next investigated the effect of texture background orientation further by calculating filling-in onset, total duration of filling-in per trial, mean filling-in episode duration and number of filling-in episodes per trial.

**Figure 3:**
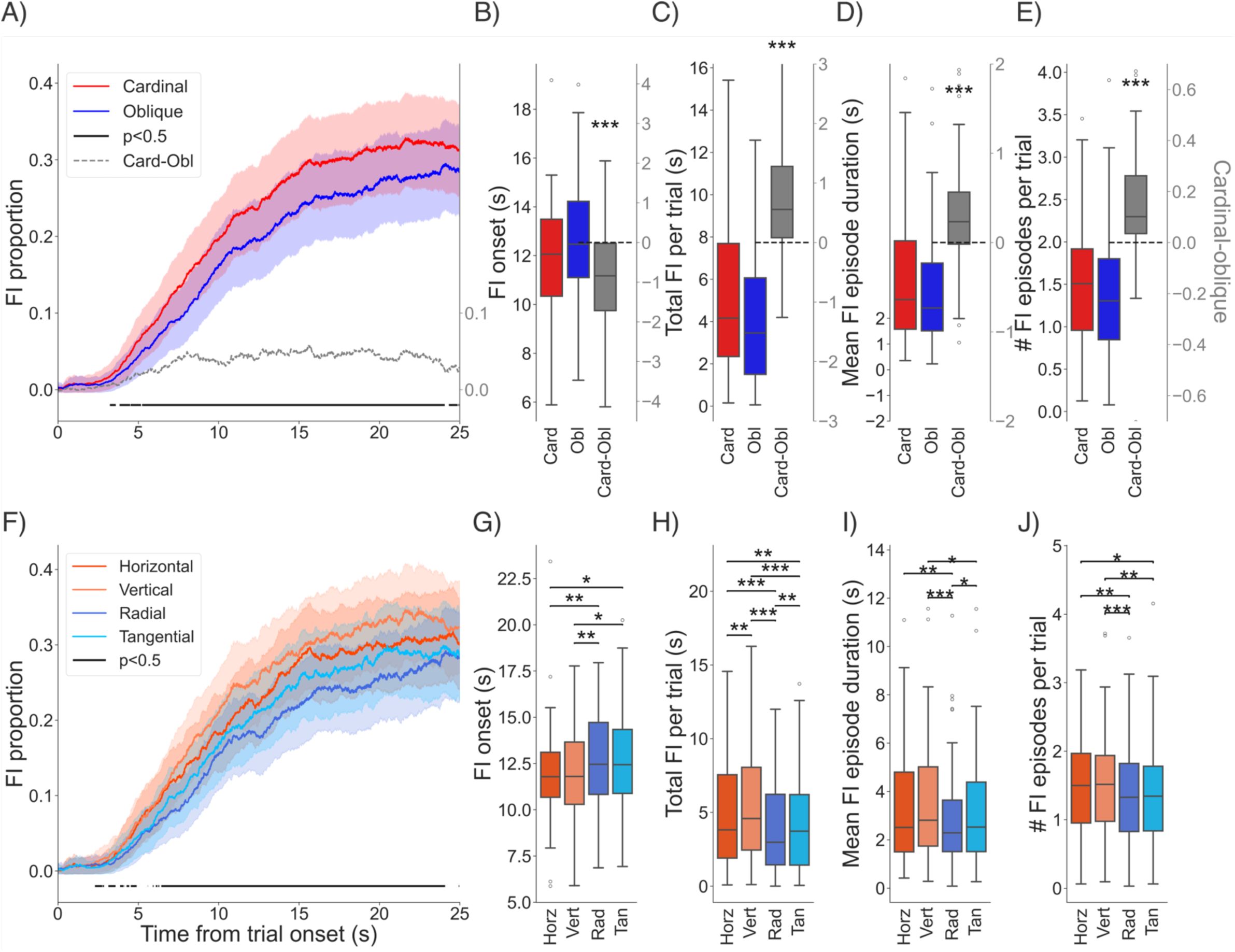
**Top row:** Cardinal (red) and oblique (blue) orientations grouped and their difference (grey). **(A)** Time course of filling-in rate. Clusters representing significant (p<0.05) difference between cardinal and oblique are indicated by the solid line at the bottom. Right axis shows difference in time course of filling-in rate between cardinal and oblique conditions. **(B)** Boxplots showing distribution of mean filling-in onset times (leftward y axis) and pairwise difference (grey, rightward axis). Stars indicate statistical difference (*<0.05 **<0.01 ***<0.001) Boxes show 25^th^, 50^th^ and 75^th^ percentile, whiskers show the rest of the distribution. Open circles show outliers. **(C)** Mean total filling-in duration per trial, conventions as in B. **(D)** Mean duration of filling-in episodes between conventions as in B. **(E)** Mean number of filling-in episodes per trial as in B. **Bottom row:** Analysis of each background orientation. **(F)** Filling-in rate across time for all 4 background orientations (dark red: horizontal, light red: vertical, dark blue: radial, light blue: tangential,). Significant clusters are indicated by the solid line at the bottom. **(G)** Boxplots of filling-in onset times. Significant differences (post-hoc tests with Wilcoxon signed-rank test) indicated by brackets along the top **(H)** Mean total filling-in duration per trial, conventions as in G. **(I)** Mean duration of filling-in episodes, conventions as in G. **(J)** Mean number of filling-in episodes per trial. Conventions as in G.

Figure 3B shows an earlier onset for the cardinal condition compared to the oblique condition. The average filling-in onset was 12.73s (*SD=*2.54) for the oblique orientations and 11.82s (*SD=*2.41) for the cardinal orientations. This difference was statistically significant with a paired t-test (*t*(57)=3.93, *p<*0.001, *d*=0.52) with a mean within-participant difference of 0.90s (*SD=*0.13). Figure 3C shows more total filling-in for cardinal than oblique texture backgrounds. The average total filling-in per trial was 5.00s (*SD=*3.71) for a cardinal background orientation and 4.15s (*SD=*3.26) for an oblique background orientation with a mean within-participant difference of 0.84s (*SD=*0.45). The difference was statistically significant (Wilcoxon signed-rank test, *W=*279, *p<*0.001, *r=*-4.46). Earlier and more total filling-in could be driven by more individual episodes of filling-in or by longer episodes (or by both), which would indicate filling-in being either more likely or more stable respectively. To investigate this, we calculated the number and length of filling-in episodes separately.

Figure 3D shows longer filling-in episodes for the cardinal condition compared to the oblique condition. The mean filling-in episode duration was 3.37s (*SD=*2.40) for cardinal background orientations and 3.04s (*SD=*2.24) for oblique background orientations, in line with a statistically significant average within-subjects difference of 0.33s (*SD=*0.16; Wilcoxon signed-rank test, *W=*342, *p<*0.001, *r=*-4.10). Figure 3E shows more filling-in episodes per trial for the cardinal condition compared to the oblique condition. There was an average count of 1.51 (*SD=*0.74) filling-in episodes per trial for cardinal background orientations and 1.38 (*SD=*0.76) for oblique background orientations. This difference was statistically significant by a Wilcoxon signed-rank test (*W=*303, *p<*0.001, *r=*-4.17) with a mean within-participant difference of 0.14 (*SD=*0.02) more episodes per trial for the cardinal orientation. Taken together, these results suggest that filling-in was indeed enhanced with a background texture of cardinal orientations, being more stable (longer lasting) and more frequent than filling-in for oblique orientations.

We next investigated whether differences existed between the two cardinal and between the two oblique orientations conditions. Previous studies have shown a tendency towards greater population response in early visual cortex to vertical compared to horizontal orientations and to radial over tangential orientations (Alink et al., 2017a; Bennett & Banks, 1991; Hansen & Essock, 2004; Hong, 2015; Leventhal & Schall, 1983; Ling et al., 2015; Maloney & Clifford, 2015; Mannion et al., 2010a, 2010b; Menceloglu et al., 2023; Rovamo et al., 1982; Sasaki et al., 2006; Schall et al., 1986; Temme et al., 1985; Westheimer, 2003, 2005). To investigate differences between the four background orientations, after combining figure orientations, we separated cardinals into vertical and horizontal. Furthermore, we grouped together the data from upper hemifield, positive-oblique and lower hemifield, negative-oblique stimuli into the condition ‘radial’, and the data from the lower hemifield, negative-oblique and upper hemifield, positive-oblique stimuli into the condition ‘tangential’. This yielded 16 trials for each of the four resulting background conditions ‘vertical’, ‘horizontal’ ‘radial’ and ‘tangential’. Figure 3F shows that the proportion of filling-in reports increased over time for all background conditions. In line with the previous analysis, this analysis suggested that there was more filling-in for the two cardinal orientations compared to the two oblique orientations. Additionally, this analysis suggested more filling-in for the vertical orientation compared to horizontal and for tangentially-oriented conditions than radially-oriented conditions. To assess the significance of the differences over time, we performed a non-parametric one-way ANOVA for repeated measures (Friedman’s test) at each timepoint, with multiple comparisons corrected for by a cluster-based permutation test. This analysis revealed clusters (black horizontal bars), indicating a statistically significant difference between the four background conditions for the majority of the trial (Figure 3F). For filling-in onset, total filling-in per trial, mean filling-in episode duration, and number of filling-in episodes per trial, we also performed a Friedman’s test (Figure 3, G-J). This analysis showed a significant main effect of background orientation on filling-in behavior for filling-in onset (#^#^=17.69, *p<*0.001; Figure 3G), total filling-in per trial (#^#^=38.21, *p<*0.001; Figure 3H), mean filling-in episode duration (#^#^=26.59, *p<*0.001; Figure 3I), and number of filling-in episodes per trial (#^#^=22.48, *p<*0.001; Figure 3J). To investigate where the differences lay, we used Wilcoxon signed-rank tests as post-hoc comparisons with Holm-Bonferroni correction. This analysis confirmed our previous results with both horizontal and vertical orientations eliciting generally earlier and more filling-in than the tangential and radial orientations. Additionally, this analysis revealed that the vertical orientation elicited the most (*M=*5.31, *SD=*3.85; Figure 3G) and earliest (*M=*11.82, *SD=*2.62; Figure 3H) filling-in, and the radial orientation the least (*M=*4.00, *SD=*3.13; Figure 3G) and latest (Figure 3H) filling-in. Post-hoc tests however revealed no statistically significant difference between vertical and horizontal orientations (pair 0°,90°) for the filling-in onset (corrected *p=*0.98; Figure 3G), mean filling-in episode duration (corrected *p=*0.14; Figure 3I) and number of filling-in episodes per trial (corrected *p=*0.30; Figure 3J). The post-hoc test for total filling-in per trial showed that vertical elicited more filling-in than horizontal (corrected *p=*0.003; Figure 3H). Additionally, post-hoc tests showed that tangential orientations elicited more filling-in (*M=*1.43, *SD=*3.36) than radial (*M=*4.00, *SD=*3.13; corrected *p=*0.006; Figure 3G) and longer episodes (*M=*3.22, *SD=*2.38) than radial (*M=*2.88, *SD=*2.19; corrected *p=*0.04; Figure 3H). Post-hoc tests indicated no statistically significant differences between radial and tangential orientations for filling-in onset (corrected *p=*0.98; Figure 3G) and number of filling-in episodes per trial (corrected *p=*0.88; Figure 3J). Taken together, these results confirmed that filling-in was modulated by the orientation of the line elements in the background texture. We observed a robust facilitation of filling-in for cardinal orientations compared to obliques in all measured parameters, and some evidence for smaller and less consistent advantages in filling-in for vertical over horizontal conditions (1 of 4 parameters) and for tangential over radial conditions (2 of 4 parameters).

### A weaker boundary representation elicits more filling-in

Previous studies have shown a greater neural response in early visual cortex to stimuli with cross-oriented surrounds than to stimuli with iso-oriented surrounds (Henry et al., 2013; Sakai & Nishimura, 2006; Self et al., 2014; Shushruth et al., 2013; Sillito et al., 1995). In our study, this effect could lead to a stronger response along the figure boundary that was orthogonal to the background orientation than the boundary that was iso-oriented to the background orientation. We used vertical and horizontal rectangular figures, the long boundary of which was three times the length of the short boundary, such that the total response to the figure boundary would be dominated by the long boundary, which were either cross-oriented or iso-oriented with respect to the vertical and horizontal background orientations. Our analysis focused exclusively on vertical and horizontal background orientations. This design choice ensured that any observed differences in boundary strength could be attributed to the iso-versus cross-oriented comparison specifically, rather than to the additional differences in cardinal versus oblique background orientations. This yielded 32 trials per condition. We expected that these configurations would yield a response to the figure boundary that was respectively stronger or weaker for the cross-oriented and iso-oriented conditions. We assessed whether modulating the expected strength of the neural response to the boundary of the figure would modulate filling-in.

Figure 4A shows that the iso-oriented conditions elicited more filling-in over time than the cross-oriented conditions although there were several overlaps between the two curves across time. This was confirmed by a cluster-based permutation test, which revealed several clusters of significant difference between the conditions.

**Figure 4:**
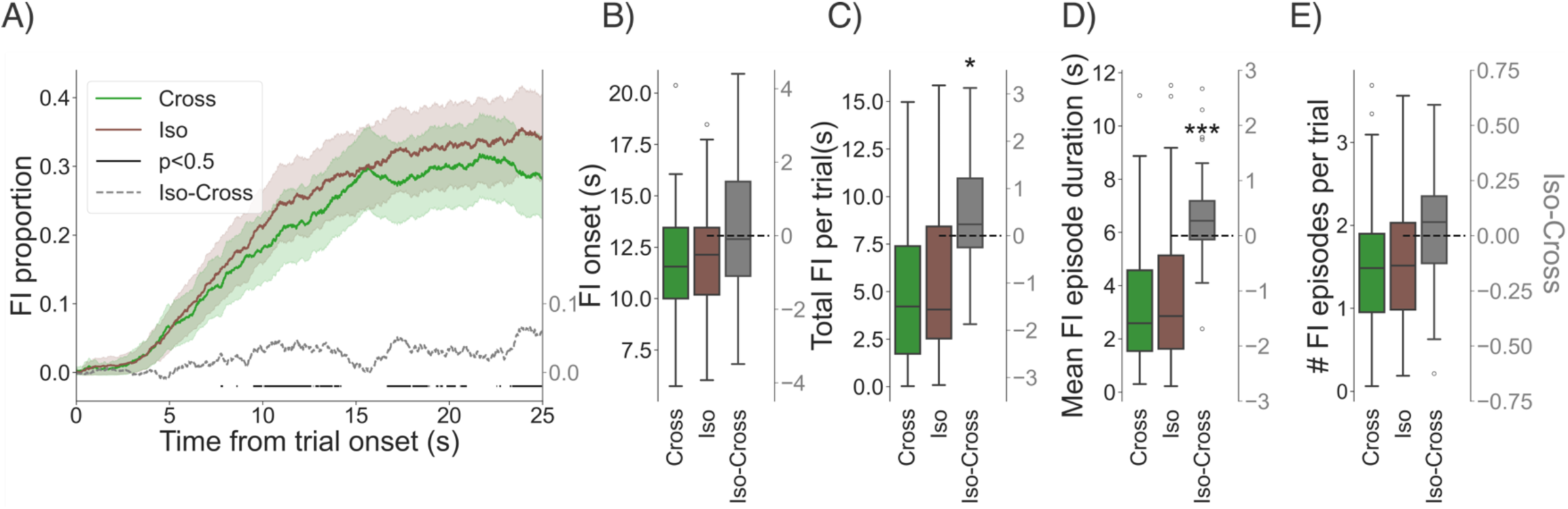
Cross-oriented (green) and iso-oriented (brown) stimuli grouped and their difference (grey). **(A)** Time course of filling-in rate. Clusters representing significant (p<0.05) difference between cross and iso-oriented stimuli are indicated by the horizontal lines at the bottom. Right axis shows difference in time course of filling-in rate between cardinal and oblique conditions**. (B)** Boxplots showing distribution of mean filling-in onset times (leftward y axis) and pairwise difference (grey, rightward axis). Stars indicate statistical difference (*<0.05 **<0.01 ***<0.001) Boxes show 25^th^, 50^th^ and 75^th^ percentile, whiskers show the rest of the distribution. Open circles show outliers. **(C)** Mean total filling-in duration per trial, conventions as in B. **(D)** Mean duration of filling-in episodes between conventions as in B. **(E)** Mean number of filling-in episodes per trial as in B.

Figure 4B shows no significant difference in filling-in onset time between the two conditions (mean onset time for cross-oriented stimuli = 11.75s *SD=*2.60; 11.87s *SD=*2.59 for iso-oriented stimuli; Students’ T-test: *t(57)*=0.50, *p=*0.62). However, we found that iso-oriented stimuli elicited more total filling-in (M=5.27s, *SD=*3.96; Figure 4C) than cross-oriented stimuli (M=4.71s, *SD=*3.53; Figure 4C). The mean within-participant difference was of 0.56s (*SD=*0.43) and was statistically significant (Wilcoxon signed-rank test, *W=*492, *r=*-2.97, *p=*0.003). Similarly, we found that iso-oriented stimuli elicited significantly longer filling-in episodes than cross-oriented stimuli, with an average filling-in episode duration of 3.54s (*SD=*2.55) for an iso-oriented stimuli and 3.09s (*SD=*2.25) for cross-oriented stimuli (Wilcoxon signed-rank test, *W=*393, *r=*-3.58, *p<*0.001; Figure 4D). We found no significant difference in the number of filling-in episodes per trial with 1.50 filling-in episodes per trial (*SD=*0.75) for iso-oriented stimuli and 1.40 (*SD=*0.78) for cross-oriented stimuli (Wilcoxon signed-rank test, *W=*715, *p=*0.38, *r=*-0.88; Figure 4E). Taken together, these results suggest that filling-in was enhanced for figures with the major axis iso-oriented with the background texture, being more stable (longer lasting) than filling-in for cross-oriented figures.

### Microsaccades rate analysis

The previous analysis showed differences in the amount of filling-in experienced for different stimulus configures which were in line with our hypothesis of filling-in being mediated by a competition between the figure boundary and background representation. However, since filling-in has been found to depend on the absence (or reduction) of microsaccades (Troncoso et al., 2008), we considered the possibility that the differences we observed could rather reflect differences in microsaccade rates between the stimulus conditions. That is, any difference in microsaccade rate between conditions would constitute a confounding factor and a potential alternative account for the pattern of results we observed.

To test for this possibility, we first calculated the microsaccade rate across the trial and compared these time courses between conditions. Figure 5A shows a relatively stable microsaccade rate of 0.26 MS/Sec across the entire length of the trial, following a brief peak in microsaccades at the trial start for both cardinal (red) and oblique (blue) background conditions. This microsaccade rate was in the same range as the microsaccade rate we have reported previously for paradigms with long trial durations (e.g. Roberts et al., 2019, 0.55MS/Sec, *SD=*0.29, during the middle of the trial). The mean microsaccade rate did not show statistical evidence of significant differences between cardinal and oblique conditions (cardinal 0.31 MS/Sec (*SD=*0.17) oblique 0.31MS/Sec (*SD=*0.17); Students’ T-test: t*(57)*=0.08, *p=*0.94). Moreover, the direction of the difference was not consistent (see difference time course, Figure 5A grey), whereas the rate of filling-in was consistently higher for the cardinal condition (Figure 2A). Similar analysis of the microsaccade rate for iso- and cross-oriented conditions (Figure 5B) also revealed no consistent statistical evidence of difference in microsaccade rate (iso-0.31 MS/Sec (*SD=*0.17) cross-0.31 MS/Sec (*SD=*0.17); Students’ T-test: *t(57)*=0.48, *p=*0.63).

**Figure 5:**
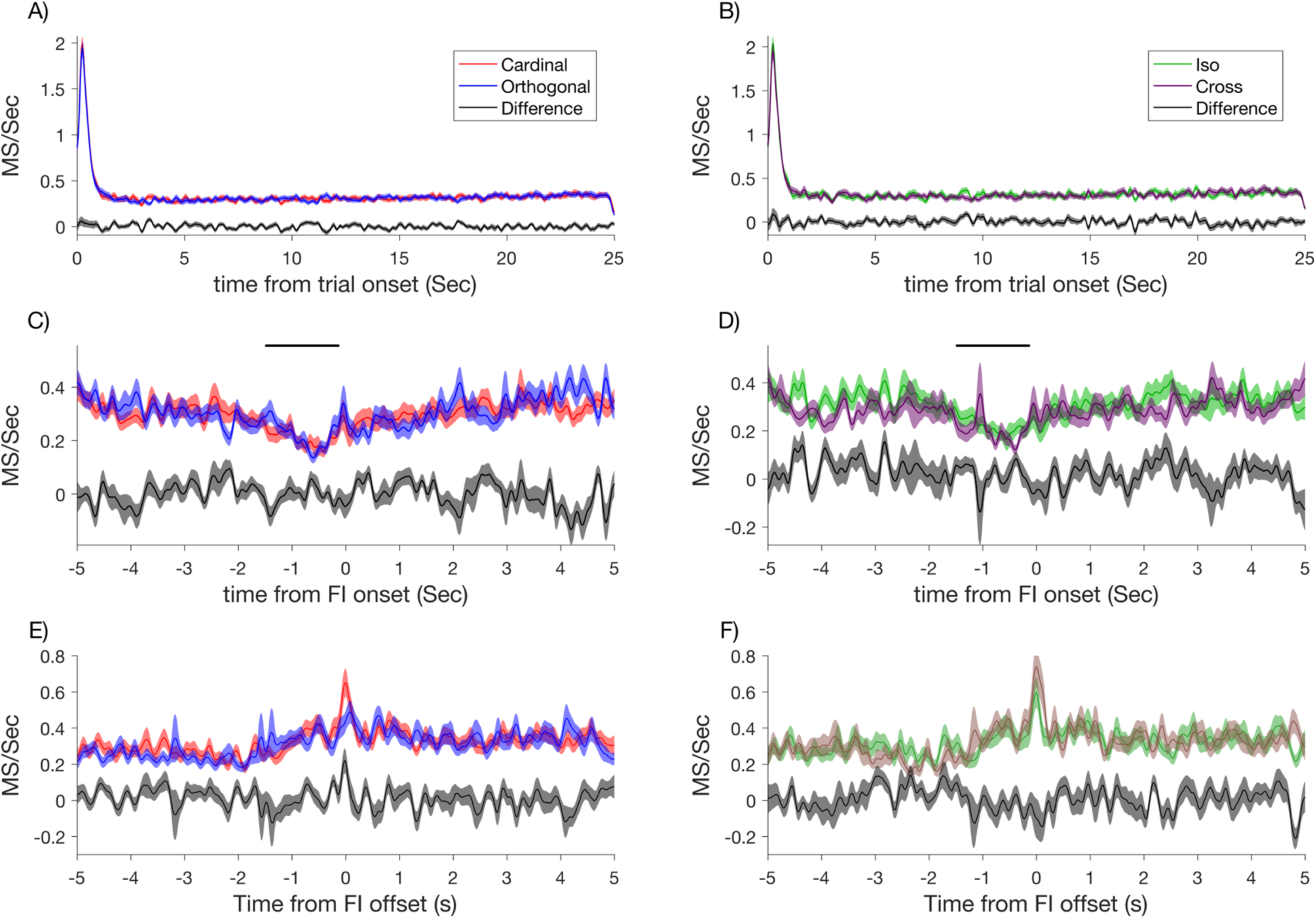
Microsaccade rate locked onto trial onset **(A, B)**, filling-in onset **(C, D)** and offset **(E, F)** for cardinal (red) and oblique (blue) and iso- (green) and cross-oriented (brown) stimuli. Difference between conditions is indicated in black. **(A, B)** Microsaccade rate across time, locked to trial onset. No significant differences between conditions were found. **(C, D)** Microsaccade rate across time, locked to filling-in onset. No significant differences between conditions were found. Time points showing a significant (cluster based permutation test, p<0.05) difference between microsaccade rate, averaged across conditions, and a null distribution (obtained by shuffled filling-in times among trials) are indicated by the horizontal bars on top of the figure panels, supporting a decrease in microsaccade rate prior to filling-in (E, F) Microsaccade rate across time, locked onto filling-in offset. There were no significant differences in microsaccade rate between paired conditions. The trend towards increasing microsaccade rate preceding filling-in offset was not significant. Conventions analogous to those in (C, D).

To further test for the involvement of microsaccades in our experiment, we calculated the microsaccade rate centered around filling-in episodes. Figure 5C shows a decrease in microsaccade rate immediately before filling-in onset for both cardinal and oblique conditions. A cluster-based statistical comparison with a permuted null distribution (see methods) revealed that this drop was statistically significant (black bar), in line with previous reports (Troncoso et al., 2008). Time-resolved comparison between conditions did not reveal clusters of statistical difference between conditions. Similar analysis of the microsaccade rate around filling-in onsets for iso- and cross-oriented conditions (Figure 5D) also revealed no consistent difference in microsaccade rate.

We repeated the analysis of microsaccade rate for filling-in offsets while again comparing cardinal to oblique conditions as well as iso- to cross-oriented conditions. Despite observing a trend towards an increase in microsaccade rate in all conditions before filling-in offset, this increase was not statistically significant after correction (Figure 5E-F). Additionally, we did not find a statistical difference in microsaccade rate between either of the paired conditions (cardinal and oblique, iso- and cross-oriented).

Next, we explored the possibility of differences in the distribution of microsaccade direction between conditions (Figure S1), since microsaccade directions have previously been reported to be influenced by the orientation of texture (Hicheur et al., 2013; Spotorno et al., 2016; Wismeijer & Gegenfurtner, 2012). We analyzed the binned distribution of all individual microsaccade directions aggregated across trials within each condition. A sign test on the mean angular difference between the conditions did not reveal significant differences for either the cardinal and oblique contrast (*p*=0.79, *BF*_10_= 0.15, mean difference = 0.19°) or the iso and cross-oriented contrast (*p*=0.60, *BF*_10_=0.18, mean difference = 4.19°) with a Bayes Factor indicating moderate evidence for the null hypothesis. Additionally, no significant differences in the mean resultant vector length of the microsaccade direction vector were observed between conditions with a Bayes Factor indicating moderate evidence for the null hypothesis (Wilcoxon signed-rank test cardinal vs oblique: *z*=0, *p*=0.34, *BF*_10_=0.22, mean difference < 0.001; iso- vs cross-oriented: *z*=0, *p*=0.45, *BF*_10_=0.15, mean difference < 0.001), meaning that there was not change in the width of the distribution of microsaccade directions between conditions. Taken together, these results indicate that the behavioral differences we observed between cardinal and oblique orientations, and between iso-oriented and cross-oriented stimuli, were driven by cortical processing rather than differences in eye movement behavior.

Finally, we evaluated whether differences in gaze position between stimulus conditions could account for the observed differences in filling-in. Systematic differences in gaze position between conditions might have altered the size of the cortical representation of the figure and thereby might have modulated filling-in (De Weerd et al., 1998). To examine this possibility, we computed the distance of gaze position from the fixation spot, and from the figure. In our analysis of gaze position relative to the fixation spot, Figure S2A shows the distribution of gaze position across all participants, with the two groups (fixation top quadrant or fixation bottom quadrant) clearly visible. Pooled across groups, we found that 95% of gaze samples fell within a radius of 1.88° of the fixation spot indicating that participants reliably maintained their gaze on the fixation spot throughout the session (Figure S2B). In our analysis of gaze position relative to the figure, we found no difference in the distance between gaze position and the figure’s center between the cardinal (mean distance = 14.67°, *SD=*1.09) and oblique (*M*=14.65°, *SD=*1.06) conditions (Wilcoxon signed-rank test *z*=1.74, *p*=0.08, *BF*_10_=0.21), and no difference between iso- (*M*=14.67, *SD=*1.09) and cross-(*M*=14.67, *SD=*1.09) conditions (Wilcoxon signed-rank test *z*=-0.44, *p*=0.66, *BF*_10_=0.15) with a Bayes Factor indicating moderate evidence for the null hypothesis. Together, these results indicate that the behavioral differences in filling-in between the cardinal and oblique conditions and the iso- and cross-oriented conditions were not explained by stimulus-driven fixation biases.

## Discussion

In a Troxler fading paradigm, where an iso-luminant blank figure overlays a dynamic texture, perceptual filling-in occurs after a period of sustained fixation and reduced microsaccade rate. Models of this effect describe a competition between separate processes that represent the background and the figure boundary, processed by respectively a boundary system and a feature system in Grossberg’s model (Grossberg, 2003). Filling-in may occur in a two-step process in which adaptation weakens boundary representation and is followed by the spreading of background information into the figure region (Gerrits et al., 1966; Gerrits & Vendrik, 1970; Gove et al., 1995; Spillmann & De Weerd, 2003). This model suggests a competition between the boundary and background representations, with filling-in occurring when the background representation is sufficiently strong to overcome the weakening boundary. To test this proposal, we investigated how varying the expected strengths of the response to the background texture and of the response to the figure boundary would modulate filling-in behavior. To this end, we manipulated the stimulus properties (background and figure orientation) in accordance with early visual cortex functional properties, and recorded reports of filling-in episodes. We hypothesized that filling-in would be most prevalent for stimulus configurations that supported a strong background response and a weak boundary response.

To vary the expected neural response to the background texture, we took advantage of the oblique effect, whereby we expected a stronger response with cardinal elements in the background than with oblique elements. In line with our expectation, we found that cardinal orientations eliciting more and longer filling-in episodes in comparison to oblique orientations. However, the oblique effect is strongest at smaller eccentricities, and weakens or even reverses at larger eccentricities, depending on the task and stimuli used (Berkley et al., 1975; Vandenbussche et al., 1986). Therefore, at the eccentricity of the figure in our stimuli it can be debated whether the oblique effect would still be very strong. This raises the possibility that the anisotropic effect of the texture background orientation on filling-in was inherited mainly from the more central parts of the texture. Future research, in which a figure presented on a spatially restricted background is placed at different distances from fixation would be required to study the relationship between the oblique effect and filling-in as a function of eccentricity.

When analyzing the four background orientations separately, we found differences between the two cardinal and between the two oblique orientations. Vertical orientations elicited more total filling-in than the horizontal orientations, in line with our expectation (Alink et al., 2017a; Freeman et al., 2011, 2013; Hansen & Essock, 2004; Maloney & Clifford, 2015; Mannion et al., 2010b, 2010a; Yacoub et al., 2008). In addition, tangential oblique orientations elicited more filling-in, with longer filling-in episodes, than radial oblique orientations. The latter finding went against our expectations since a majority of studies suggest a greater population response to radial over tangential orientations (Alink et al., 2017a; Bennett & Banks, 1991; Freeman et al., 2013; Hong, 2015; Leventhal & Schall, 1983; Levick & Thibos, 1982; Ling et al., 2015; Maloney & Clifford, 2015; Mannion et al., 2010a, 2010b; Menceloglu et al., 2023; Rovamo et al., 1982; Sasaki et al., 2006; Schall et al., 1986; Temme et al., 1985; Westheimer, 2003, 2005). However, not all studies have consistently reported the radial over tangential advantage (Alink et al., 2017b; Champion & Warren, 2017; Mannion et al., 2010b).

To vary the expected response to the figure boundary, we took advantage of the orientation specificity of surround modulation effects known from V1 neurons, whereby suppression from the surround is stronger when surround elements are iso-oriented to a central stimulus than when they are cross-oriented. Thus, we manipulated the orientation difference between the long axis of the rectangular figure and the orientation of the background texture. We found more filling-in when the long boundary of the figure was iso-oriented with the background than when it was cross-oriented.

Although our data confirmed the expectation of stronger filling-in in the iso-oriented condition, the stronger filling-in in that condition could also be explained by an alternative theory. Some studies have investigated a modulation of perceptual filling-in depending on figure-background boundary saliency (De Weerd et al., 1998; Gyoba, 1997; Stürzel & Spillmann, 2001; Welchman & Harris, 2001). These studies found that a figure with a higher perceptual saliency delayed perceptual filling-in compared to a less salient figure. In our stimulus, the rectangular figure with its long axis iso-oriented with the line elements entailed that only the short edges were reinforced by the orientation contrast between line element and figure edge, whereas the opposite was true when the long axis of the figure was cross-oriented to the orientation of the line elements. We hypothesized that the strength of the boundary response would be modulated according to the orientation tuning of surround suppression in early visual areas. However, orientation contrast may also represent occlusion cues, making the figure ‘pop-out’ from the background and so modulating the perceptual saliency of the figure. Thus, the reduced filling-in observed for cross-oriented figures in our study could represent the figure being more salient, rather than (or in addition to) representing the operation of center-surround mechanisms as we have described. Crucially, while both alternative interpretations rely on orientation contrast, the saliency mechanism implies the involvement of additional higher-order cognitive processes. While our data do not preclude the involvement of such processes, neither do they necessarily support them.

Our study is the first to our knowledge to directly compare filling-in between the upper and lower hemifields. Previous studies have presented the filling-in figure either in the upper (e.g., Ramachandran & Gregory, 1991; Sakaguchi, 2001; Shimojo et al., 2001), or the lower (e.g., De Weerd et al., 1995, 1998) hemifield, and studies that presented the figure in both hemifields (e.g., Meng et al., 2005; Spillmann & Kurtenbach, 1992; Troncoso et al., 2008) did not analyze the difference in filling-in between hemifields. Many studies have shown a better visual performance in the lower hemifield compared to the upper hemifield in a variety of discrimination and detection tasks including orientation discrimination (e.g., Barbot et al., 2021; Carrasco et al., 2001), line bisection (e.g., Nielsen et al., 1999) and visually guided pointing tasks (Goodale & Danckert, 2001). Furthermore, the strength of subjective contours was stronger lower compared to upper hemifields (Rubin et al., 1996; Shapley et al., 2003). Based on this finding, one might expect a difference in filling-in, but we did not observe a difference in filling-in behavior between presentation in the upper or the lower hemifield. The main difference between our stimulus and these studies is that they used a central figure subtending a limited area while we used stimuli in the periphery of the visual field, in a location possibly less affected by this asymmetry found for illusory contours.

Our study included the largest sample yet in a filling-in study to our knowledge, 58 participants compared to 27 in the next largest sample (Ehinger et al., 2017). We took advantage of the size of the sample to characterize the central tendency of filling-in behavior in this sample. We did not observe a multi-modal distribution of participants in filling-in behavior, suggesting that participants cannot be categorized as some experiencing “a lot of” and others experiencing “no” filling-in, and this was true for all extracted parameters of filling-in, including, average filling-in onset, duration of filling-in episodes and count of filling-in episodes per trial. Interestingly, in our large sample all participants reported experiencing some degree of perceptual filling-in, supporting perceptual filling-in as a robust and general phenomenon in the population.

Several previous studies have validated the robustness of perceptual filling-in by demonstrating an alignment between subjective participant reports to the illusion and catch-trial performance across multiple forms of perceptual filling-in (Ehinger et al., 2017; Levinson et al., 2025; Welchman & Harris, 2001b). Although in the present study, we did not use a catch-trial approach, the consistency of responses to filling-in and physical lures in these studies provides indirect support for the robustness and validity of subjective reports as a measure of filling-in also in our paradigm.

Changes in microsaccade rate modulate perceptual filling-in (Troncoso et al., 2008), and therefore potential differences in microsaccade rate among conditions could have contributed to differences in perceptual filling-in. We therefore analyzed microsaccade rate dynamics around filling-in events in the different conditions. In our experiments, stimulus configurations did not modulate microsaccade rate, either throughout the trial, or around filling-in onset and filling-in offset. However, across all conditions we found that microsaccade rate decreased before filling-in onset, in line with Troncoso et al.’s, (2008) findings. They also reported an increase in microsaccade rate prior to filling-in offset. We found a similar trend in our data, which, however, was not statistically significant. Next, we analyzed microsaccade directions, following evidence that texture element orientation modulates microsaccade direction (Hicheur et al., 2013; Spotorno et al., 2016; Wismeijer & Gegenfurtner, 2012). Using circular statistics, we found no statistically significant differences in microsaccade direction among conditions, and no differences in resultant microsaccade mean direction vector length. Given that microsaccade rate, and direction were not significantly different between conditions, we conclude that the differences in filling-in behavior among conditions were not explained by differences in microsaccade behavior between the different stimulus conditions. Finally, we analyzed gaze positions to determine whether stimulus properties systematically influenced where participants maintained their gaze. Changes in gaze position relative to the figure would influence the size of its representation in retinotopic cortex, which is a factor known to influence perceptual filling-in (De Weerd et al., 1998). We found no evidence, however, that gaze position varied with stimulus condition. We therefore conclude that that the differences in filling-in behavior were also not explained by differences in gaze position.

Our results and the conceptual model we used to interpret them are qualitatively related to existing, more detailed theoretical frameworks (Grossberg, 2016; Grossberg & Mingolla, 1985b, 1985a; Grossberg & Pessoa, 1998; Neumann et al., 2001; Peters et al., 2010). Future computational work may test to what extent our results would fit quantitative predictions from detailed computational models of surface perception, and to what extent our results might require amending some aspects of these models. In addition, future neuroimaging studies could replicate our stimulus manipulation to test their expected effects on the strength of boundary and texture responses, and on perceptual filling-in. In addition, the hypothesis that early visual cortex contributes to filling-in, which is supported by our data, could be tested by applying suppressive transcranial magnetic stimulation protocols to early visual cortex in both behavioral and neuroimaging experiments. Furthermore, in future studies, the approach in which perceptual filling-in is tested as a function of stimulus manipulations modulating neuronal responses in early visual cortex could be expanded to test contributions of higher-level cortical areas.

## Conclusions

Our study shows that stimuli expected to elicit a strong texture background representation and a weak boundary representation led to more filling-in, with more and longer filling-in episodes. These results indicate that the two stages in the two-step model of filling-in, boundary adaptation and filling-in, are influenced by early visual cortical mechanisms.

