## Supplementary Figure 1 and 2 for "Stimulus dependent modulation of perceptual filling-in is predicted by the properties of early visual cortex"

**Supplementary Materials**

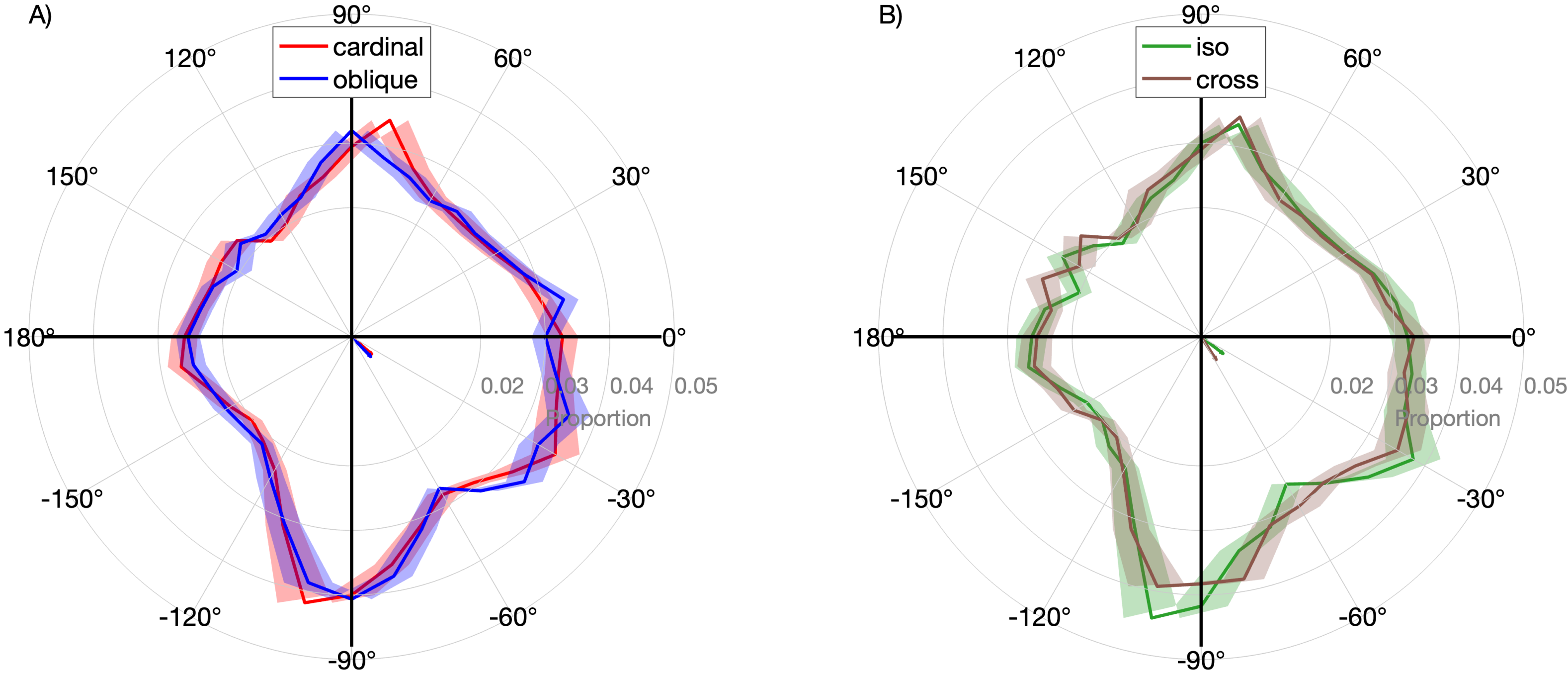


*Figure S1: Proportion of microsaccade direction for:* ***A)*** *cardinal (red) and oblique (blue),* ***B)*** *iso- (green) and cross- (brown) conditions. Shaded areas indicate standard error of the mean. The small lines at the center of the polar plots represent the mean vector for each condition. No differences in microsaccade direction or mean vector length were found between conditions (Sign test on pairwise differences and Wilcoxon signed-rank test, both p > .05).*

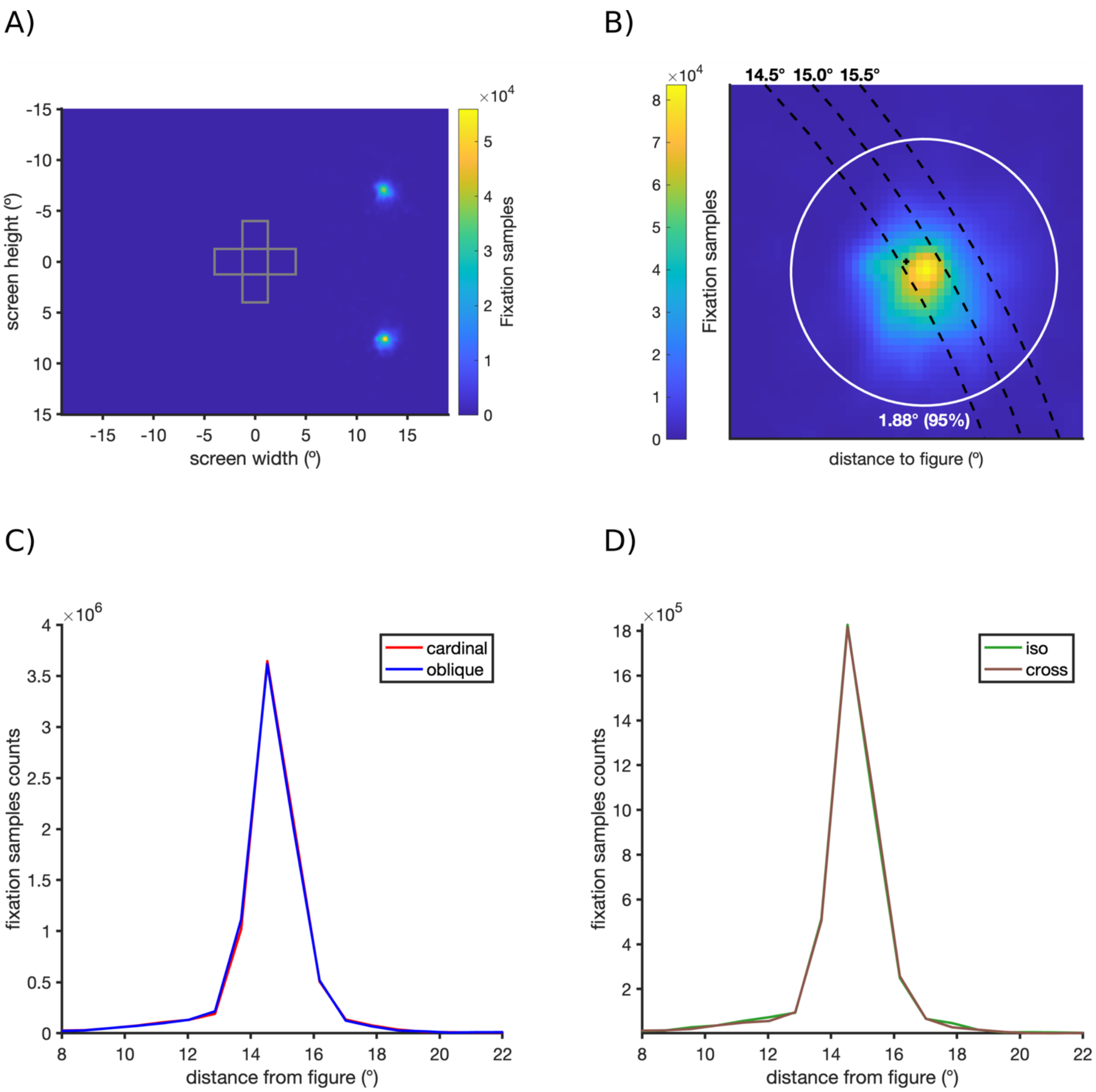


*Figure S2: Gaze position location relative to the figure location and physical fixation spot **(A)** Gaze position density map across the entire visual field. **(B)** gaze position density map at fixation spot with data from upper fixation location and lower fixation location combined (Y values of the lower fixation location group have been mirrored). Black cross indicates the fixation spot location. Dashed contours indicate distance from the figure center. Red circle indicates the contour containing 95% of all gaze locations. **(C, D)** Distributions of gaze location distance to the figure center for cardinal (red), oblique (blue), iso- (green) and cross- (brown) oriented stimuli. No difference in fixation distance to the figure location (Wilcoxon signed-rank test, all p>0.05).*
